# A novel bacteriophage with broad host-range against *Clostridioides difficile* ribotype 078 elucidates the phage receptor

**DOI:** 10.1101/2021.05.26.445907

**Authors:** M.J Whittle, T.W Bilverstone, R.J van Esveld, A.C. Lücke, M.M Lister, S.A Kuehne, N.P Minton

**Affiliations:** Clostridia Research Group, BBSRC/EPSRC Synthetic Biology Research Centre (SBRC), School of Life Sciences, Biodiscovery Institute, The University of Nottingham, Nottingham, NG7 2RD, UK; NIHR Nottingham Biomedical Research Centre, Nottingham University Hospitals NHS Trust and the University of Nottingham, Nottingham, NG7 2RD, UK; Faculty of Medicine, Leiden University Medical Centre, the Netherlands; Hannover Medical School, Carl-Neuberg-Straße 1, 30625 Hannover, Germany; Department of Clinical Microbiology, Queen’s Medical Centre, Nottingham University Hospitals NHS Trust, Nottingham, UK, UK; Oral Microbiology Group, School of Dentistry and Institute of Microbiology and Infection, College of Medical and Dental Sciences, The University of Birmingham, Birmingham, B5 7EG, UK

**Keywords:** Bacteriophage, phage therapy, *Clostridioides difficile (Clostridium difficile)*, S-layer, SlpA

## Abstract

Bacteriophage represent a promising option for the treatment of *Clostridioides difficile* (formerly *Clostridium difficile*) infection (CDI), which at present relies on conventional antibiotic therapy. The specificity of bacteriophages should prevent the dysbiosis of the colonic microbiota associated with the treatment of CDI with antibiotics. Whilst numerous phages have been isolated, none have been characterised with broad host-range activity towards PCR ribotype (RT) 078 *C. difficile* strains despite their considerable relevance to medicine and agriculture. In this study, we isolated four novel *C. difficile* Myoviruses: ΦCD08011, ΦCD418, ΦCD1801 and ΦCD2301. Their characterisation revealed that each was comparable with other *C. difficile* phages described in the literature, with the exception of ΦCD1801 which exhibited a broad host-range activity towards RT 078, infecting 15/16 (93.8%) of the clinical isolates tested. In order for wild-type phages to be exploited in the effective treatment of CDI, an optimal phage cocktail must be assembled that provides broad coverage against all *C. difficile* RTs. In an attempt to advance these efforts, we conducted a series of fundamental experiments that identified the *C. difficile* SlpA, the major constituent of the *C. difficile* surface-layer (S-layer), as the phage receptor. Thus, we demonstrated that ΦCD1801 could only bind to RT 012 or RT 027 strains in the presence of a plasmid-borne S-layer cassette corresponding to RT 078. Armed with this information, efforts should now be directed towards the isolation of phages with broad host-range activity against each of the fourteen described S-layer cassette types which could form the basis of an effective cocktail active against a wide range of *C. difficile* isolates.

**Importance:** Research into phage therapy has seen a resurgence in recent years owing to growing concerns regarding antimicrobial resistance. Phage research for potential therapy against *Clostridium difficile* infection (CDI) is in its infancy, where an optimal “one size fits all” phage cocktail is yet to be derived. The pursuit thus far, has aimed to find phages with the broadest possible host-range. Although, for *C. difficile* strains belonging to certain PCR ribotypes (RTs), in particular RT 078, phages with broad-host range activity are yet to be discovered. In this study, we isolate 4 novel Myoviruses including ΦCD1801, which exerts the broadest host-range activity towards RT 078 reported in the literature. Through the application of ΦCD1801 to robust binding assays, we elucidate SlpA as the phage receptor on the bacterial cell surface. Our finding suggests that an optimal “one size fits all” combinatorial phage cocktail, could theoretically comprise 14 phages, each targeting one of the 14 described S-layer cassettes of *C. difficile*.

## Introduction

*Clostridioides difficile* (formerly *Clostridium difficile* [1]) is the leading cause of hospital-associated diarrhoea in the developed world, responsible for up to 29,000 deaths per annum in the USA [2]. *C. difficile* infection (CDI) ensues from dysbiosis of the gut microbiota, in response to broad-spectrum antibiotic treatment [3]. Up to 65% of patients suffer recurrent infection or relapse, following treatment of CDI with metronidazole or vancomycin [4]. This phenomenon is a consequence of the spore-forming nature of *C. difficile*, concomitant with the reduced-diversity microbiota following sustained antibiotic therapy [5]. This unfortunate chain of circumstance, whereby the antibiotic for the treatment of CDI, is also the predisposing risk factor for its contraction, calls for a more directed approach to combatting this infection. An approach that minimally disrupts the diversity of the gut microbiota.

Bacteriophages (phages) are generally considered as narrow host-range viruses, where host specificity can be observed at the genus, species or sub-species level [6]. Consequently, bacteriophage might represent an appropriate narrow-spectrum therapy for the treatment of CDI. To-date, many phages infecting *C. difficile* have been characterised, all of which, are temperate Myoviridae/Siphoviridae belonging to the Caudovirales order of phages. However, no phage has been described with broad host–range activity towards PCR ribotype 078 (RT 078), strains of which are of considerable clinical and agricultural relevance [7].

Whilst the efficacy of single-phage therapy has little remedial effect *in vivo*, combinatorial therapy has demonstrated some merit. Therein, a cocktail of phages was able to delay the time to end-point by almost 100% in hamsters infected with one strain of *C. difficile* [8]. These promising data warrant further study into optimal phage cocktail combinations.

The efforts towards combinatorial phage cocktails would be considerably boosted, had the phage receptor on the surface of the bacterial cell, been identified. Although two research articles have suggested that the surface layer (S-layer) constituent, SlpA, represents the likely phage receptor candidate [9, 10], there exists a lack of experimental evidence to verify the crucial relationship between the infecting bacteriophage and the identity of the host S-layer.

In this study, we isolated and characterised four novel bacteriophages, one of which, ΦCD1801, possessed broad host-range activity against *C. difficile* PCR ribotype 078 (RT 078). Through the plasmid-borne expression of RT 078 S-layer cassettes in RT 012 and RT 027 strains, we were able to demonstrate cross-ribotype binding, thus affirming SlpA, as the phage receptor for *C. difficile*.

## Results and Discussion

### Isolation and visualisation of four novel Myoviruses

We sought to isolate phages with infective capacity towards RT 078. RT 078 strains are often considered potentially-hypervirulent [11]. Indeed, strains possess the binary toxin genes (*C. difficile* Transferase, CDT), whilst their clinical presentation is comparable to that of the notorious RT 027 [12]: the hypervirulent ribotype responsible for severe outbreaks across North America and Europe [13]. To enhance our efforts, we obtained a library of clinical isolates from CDI-positive patients at the Queens Medical centre (Nottingham, UK), which included eight novel RT 078 isolates (See Table S1 for novel strains described in this article). Using CD1801 as an isolation host, we were able to isolate ΦCD1801 from an anaerobic digester sample derived from the Stoke Bardolph sewage treatment plant (Nottinghamshire, UK). In parallel, we isolated ΦCD08011, ΦCD2301 and ΦCD522418 (hereafter referred to as ΦCD418) from RT 002, 014 and 023 hosts, respectively.

TEM analysis revealed that ΦCD418, ΦCD2301 and ΦCD1801 possessed contractile tails (Fig. 1a-c), suggesting they belonged to the *Caudovirales* order of tailed phages and were, like most of the published *C. difficile* phages, Myoviruses. Indeed, the tail and capsid measurements are in-line with those previously reported for *C. difficile* Myoviruses [14, 15]. The imaging results were less clear for ΦCD08011. Wherein, after multiple experiments, phage particles always appeared with contracted tails and empty capsids indicative of DNA release (Fig. 1d). In light of these issues, we are unable to definitively state that ΦCD08011 is a member of the Myoviridae family.

**Figure 1:**
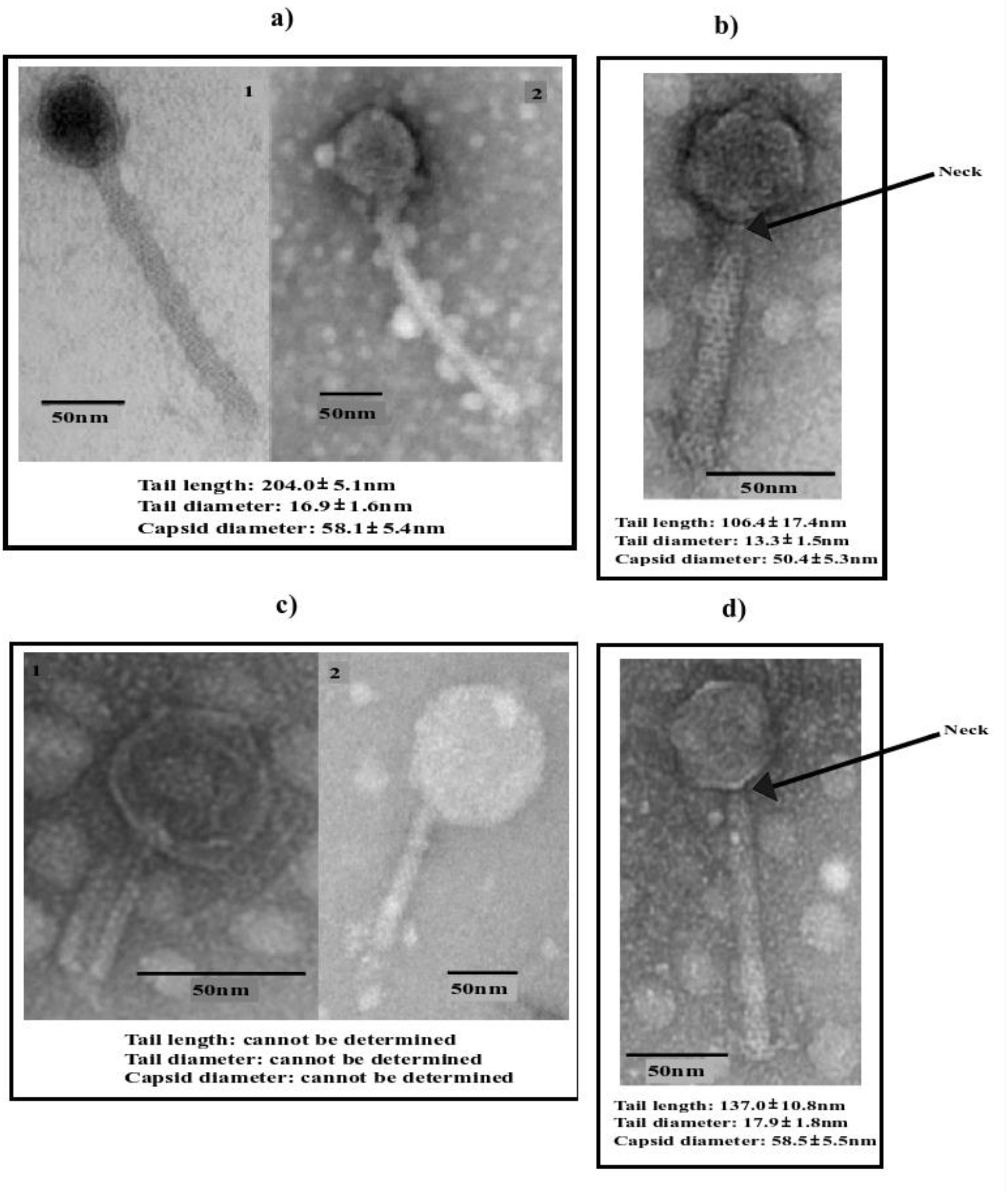
Phage particle morphology as visualised using TEM. for **a)** ΦCD418; **b)** ΦCD2301; **c)** ΦCD1801; **and d)** ΦCD08011. Measurements represent the mean ±SD of values of 5 individual phage particles.

### Phage genome sequencing, annotation and analysis

Full genome sequencing of ΦCD1801 followed by *de novo* assembly, revealed a 44,363bp circular genome with a GC content of 28.87%. Artemis software predicted the genome to contain 50 ORFs, of which putative function could be assigned to 32. A graphical representation of the ΦCD1801 genome is provided in Fig. 2a. A lytic repressor protein (CD1801_gp35) was annotated by sequence alignment with repressor proteins derived from other published *C. difficile* phage sequences using the EMBOSS pairwise sequence-alignment tool [16]. Doing so, unveiled 100% amino acid similarity with the repressor protein in *C. difficile* phage ΦCD27 [17]. A head connector protein (CD1801_gp8) was also located within the genome, further confirming the classification of ΦCD1801 as a Myovirus. The genome was closed via PCR with primers annealing to the left and right flanks of the assembled contig. Primer sequences for the closure of phage genomes are provided in Table S2.

**Figure 2:**
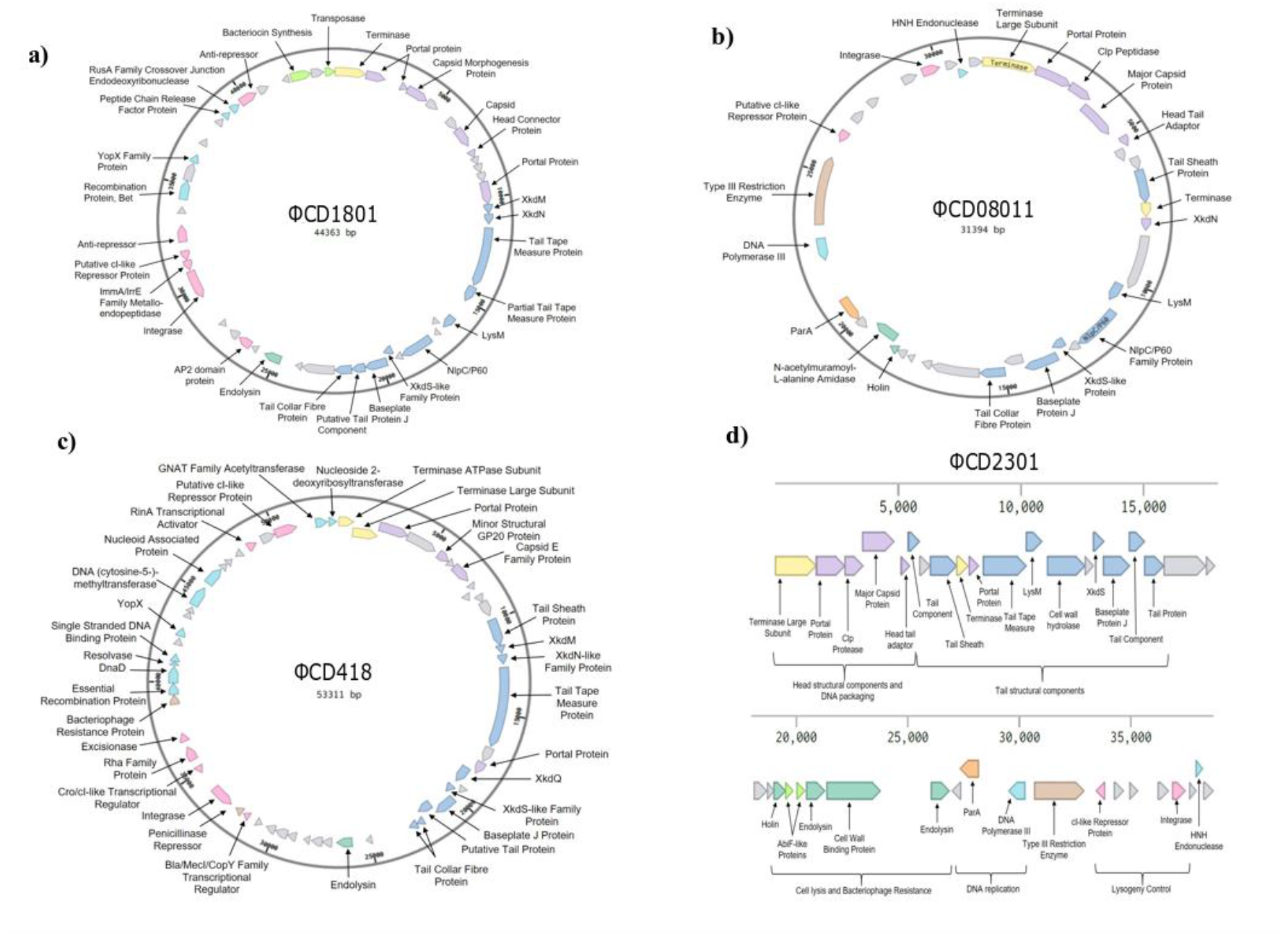
Graphical representation of phage genomes. **a)** ΦCD1801; **b)** ΦCD08011; **c)** ΦCD418**;and d)** ΦCD2301;.Assembled using CLC Genomics Workbench and manually annotated using Artemis, BLAST, UniProt and pfam. Notably, the genome contains an integrase gene indicating ΦCD1801 is a temperate phage. Proteins encoding the capsid and tail including the base plate protein were identified.

Using the abovementioned analyses, closed genome maps were generated for ΦCD08011 and ΦCD418 with genome lengths of 31,394 and 53,311bp, respectively (Fig. 2b-c). Both genomes comprised dsDNA with GC contents of 29.81% (ΦCD08011) and 29.07% (ΦCD418). A total of 35 ORFs were detected for ΦCD08011 of which 23 were assigned putative function, compared with 35 putative functional protein-coding genes, from a total of 58 for ΦCD418.

It was not possible to generate a closed genome for Φ2301. Whilst the 38,695bp dsDNA genome could be assembled into one single contig, it was not possible to close the genome via PCR at the left and right flanks of the assembled reads, despite repeated attempts. As such, we present this genome as a linear fragment (Fig. 2d). A total of 39 ORFS were detected for Φ2301 of which 27 could be assigned putative function.

Identification of integrase genes at gene position 33 (CD1801_gp33) for ΦCD08011 and ΦCD1801, indicates that, as is the case for all other reported phages infecting *C. difficile*, they are temperate. As integrase genes were identified at gp36 and gp37 for ΦCD418 and ΦCD2301 respectively. These data are confounded by the generation of lysogens for each of the four phages described herein.

*In silico* analysis revealed the presence of integrase genes in the genomes of all four phages, corresponding to CD1801_gp33, CD08011_gp33, CD418_gp36 and CD2301_gp37 for ΦCD1801, ΦCD08011, ΦCD418 and ΦCD2301, respectively. Their presence suggests that all four are, in common with all previously isolated *C. difficile* phage, temperate in nature. This conclusion was confirmed by the subsequent isolation of *C. difficile* lysogens for each phage.

Annotated phage genomes were submitted to GenBank as a BankIt submission. The accession numbers for each genome are as follows: ΦCD1801 (MW512570); ΦCD08011 (MW512572); ΦCD418 (MW512573); and Φ2301 (MW512571).

### Phage host range testing

Thus far, we had isolated four Myovirus phages that individually infect at least one RT 001, 014, 023 and 078 isolate. Genomic and phenotypic analysis thereof, suggests that the novel phages are comparable to other phages reported for *C. difficile*. Hitherto, there have been few phages characterised with infective capacity for RT 078. Most of which demonstrated very narrow host-ranges within the ribotype. For example, when a panel of seven phages were screened for their ability to infect eight RT 078 isolates, seven had no host-range coverage whatsoever, whilst one phage was able to infect 3/8 strains, representing 38% host-range activity [8].

To determine the host-range coverage of our novel phages, we adopted a standard double agar overlay plaque assay (see Methods) on 162 clinical isolates of *C. difficile* of varying RTs. This analysis revealed that ΦCD1801 had broad host-range activity towards RT 078. Infection was observed for 15/16 isolates with varying efficiencies of plating (see experimental), representing 93.8% coverage (Fig. 3). To our knowledge, these data indicate that ΦCD1801 has the greatest reported host-range coverage within RT 078. To ascertain why our phage was unable to infect the resistant strain (CD2315), we analysed the genome listing previously provided to NCBI by our research group (Accession CP068554.1). Analysis using the PHASTER web tool [18], identified the presence of an intact prophage. Further inspection by means of EMBOSS pairwise sequence analysis [17], uncovered 100% nucleotide identity between the lysogenic repressor protein of ΦCD1801 and the prophage contained within the genome of CD2315. Taken together, it is likely that the presence of this identical repressor protein is responsible for the lysogenic immunity of CD2315 towards phage infection by ΦCD1801.

**Figure 3:**
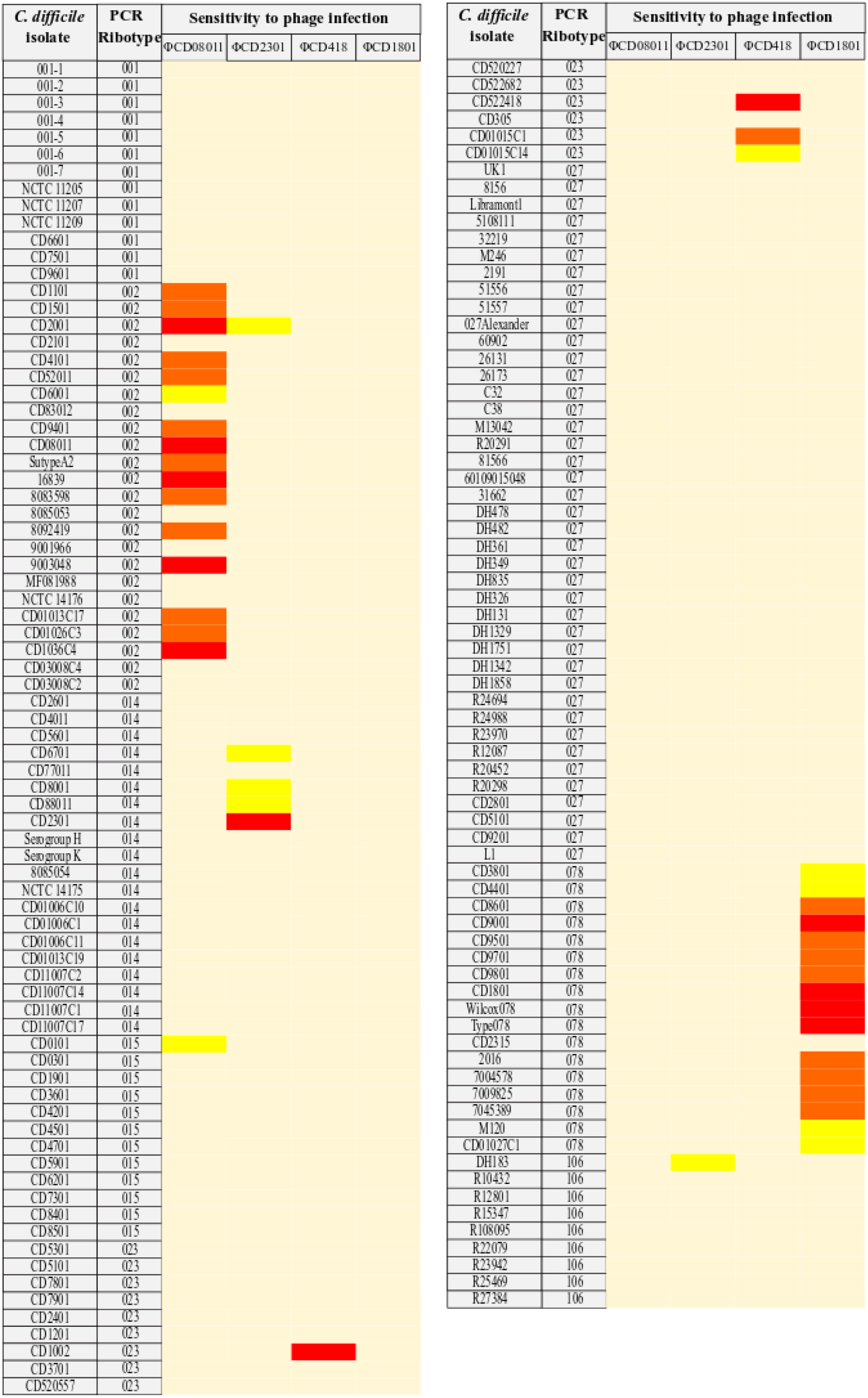
Heat map of *C. difficile* sensitivity to bacteriophage infection. Phage host range was determined using standard plaque assays for phages ΦCD08011, ΦCD2301, ΦCD418 and ΦCD1801. Efficiency of plating (EOP) values are depicted for each infection study represented by colour. Magenta: resistant strain; yellow: EOP <0.1; orange: EOP 0.1-1; red: EOP >1.

In a similar manor to the above, ΦCD08011 possessed a stringent affinity towards the RT from which it was isolated. Indeed, the phage could infect 15/23 RT 002 isolates representing 65.2% coverage (Fig. 3). The remaining two phages demonstrated remarkably lower host range activity than the abovementioned phages. ΦCD2301 was only able to infect 3/19 RT 014 strains representing only 15.8% host range coverage. In addition to the three RT 014 strains, ΦCD2301 was shown to infect the RT 106 isolate CDDH183, albeit with a low EOP value. Finally, ΦCD418 was able to infect only 3/14 of the tested RT 023 strains representing 21.4% host range coverage.

Taken together, it appears that the phages isolated herein appear to have essentially strict sensitivity to one particular RT. This differs from most of the previously published phages in which cross-ribotype sensitivity is frequently observed [15]. This phenomenon is unexplained at present but could relate to the sample origin for phage isolation. In our study, we used sewage samples from the UK which are ultimately derived from human faeces. Other studies utilised purely environmental samples for phage isolation, for example soil [15].

### Identification of the S-layer as the surface receptor for ΦCD1801

The S-layer of *C. difficile* is a para-crystalline protein that coats the entire bacterial cell, comprised of a precursor protein SlpA that is post-translationally cleaved into high molecular weight and low molecular weight SlpA derivatives [19]. The gene encoding SlpA is located within a hypervariable S-layer cassette (SLC) comprising a five-gene cluster containing *slpA, sec2A, cwp2, cwp66* and *cwp2790*. Thus far, 14 S-layer cassette types (SLCT) have been determined, where the variability is mainly ascribed to sequence differences within the low molecular weight component of *slpA* [20]. Research conducted on a novel R-type bacteriocin (Avidocin-CD), uncovered SlpA as the surface receptor for these novel antibacterial agents, since sensitivity could be conferred to a resistant strain in an SLCT-dependent manner [9]. Owing to R-type bacteriocins naturally resembling Myovirus phage tail-proteins [21], the authors provided a clear indication that SlpA is a likely surface receptor candidate for Myovirus phage infection in *C. difficile*. Although Phothichaisri and colleagues have demonstrated a physical interaction between phage particles and the surface layer of *C. difficile* by means of native-PAGE analysis [10], hitherto, no research has been conducted to verify the relationship between SLCTs and their interaction with *C. difficile* phages.

In order to determine the relationship between various SLCTs and our bacteriophage, we conjugated pJAK002 comprising pRPF185, expressing the RT 078-derived hybrid S-layer cassette (H2/6), as well as the individual H2 (pJAK023) and H6 SLCs (pJAK018) [9], into the RT 012 strain 630. Following successful conjugation, the recipient strain was assessed for its ability to bind ΦCD1801, by means of a binding assay. These analyses revealed that CD630 was unable to bind ΦCD1801 (Fig. 4). Thus, no substantial reduction in titer was observed (≥1-log), between the initial phage inoculum and the number of unbound phage particles following plaque assay in strain CD1801. However, in the presence of plasmid-borne SLCs H2/6 and 6 encoded on pJAK002 and pJAK018, respectively, ΦCD1801 bound to CD630 as indicated by a substantial decrease in the number of unbound phage particles observed (Fig. 4). The same pattern was observed when we tested the two RT 027 strains CDDH1916 and CD31662 (Fig. S1). Taken together, these data demonstrate that phage binding for ΦCD1801, is dependent on the SLCT. Therefore, our results corroborate the above-mentioned findings [9, 10], thus affirming SlpA as the surface receptor for bacteriophage infection by *C. difficile* Myoviruses.

**Figure 4:**
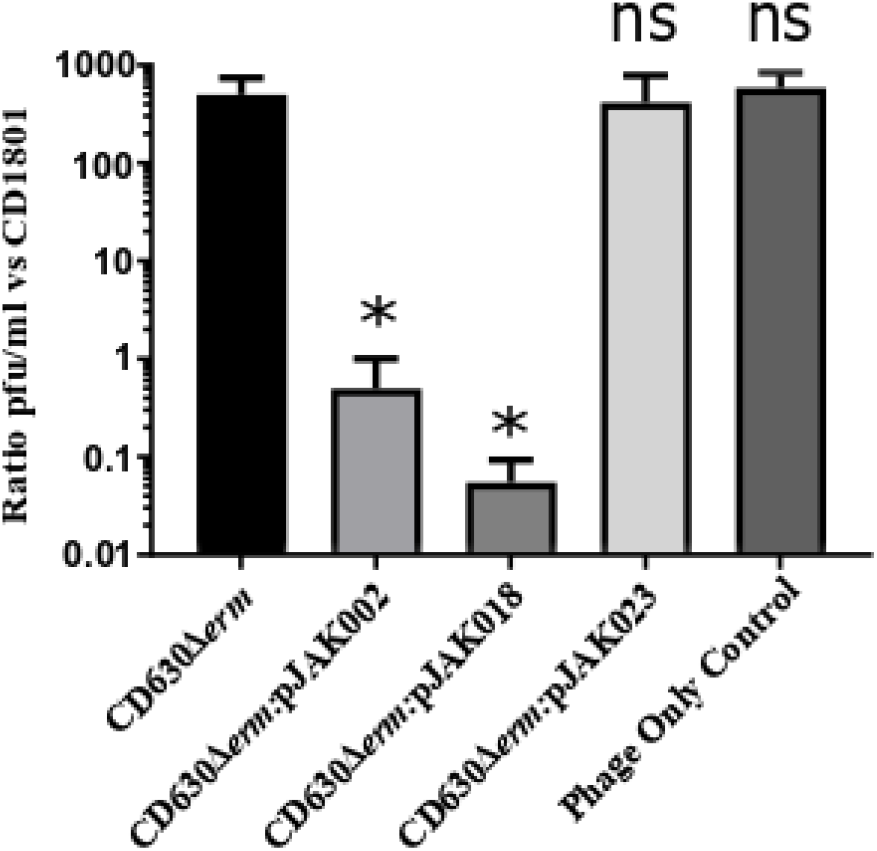
Phage binding is SLCT-dependent in RT 012. The capacity for CD630 to bind to ΦCD1801 was assessed by means of a phage binding assay with wild-type CD630 and CD630 harnessing plasmid-borne SLC H2/6 (pJK002), SLC 6 (pJK018) and SLC 2 (pJK023) under the control of a tetracycline inducible promoter. Following co-incubation with wild-type or plasmid-bearing CD630, the titer of free ΦCD1801 particles was assessed through a plaque assay using the propagating strain CD18101 as an indicator. Data represent the mean ±SD of 3 biological replicates P-* <0.05 according to One-Way ANOVA followed by Dunnett’s multiple comparison test.

Given the notion that 14 SLCTs exist for *C. difficile*, it can be reasonably assumed that an optimal phage cocktail of up-to 14 phages will be required for broad host-range coverage against CDI. To achieve this milestone, efforts must first be made towards the isolation and characterisation of phages, which are able to individually infect strains harbouring one of the 14 SLCTs. Doing so has the potential to considerably enhance the therapeutic prospect of phage therapy for the treatment of CDI.

## Conclusions

Four novel bacteriophages infecting *C. difficile* have been isolated from UK sewage samples: ΦCD08011, ΦCD418, ΦCD1801 and ΦCD2301. While ΦCD08011, ΦCD418 ΦCD1801 and ΦCD2301 were comparable to other phages reported in the literature, ΦCD1801 was shown to possess the broadest reported host-range activity towards RT 078 strains of *C. difficile*, wherein 15/16 (93.75%) of clinical isolates were susceptible to lysogenic infection. To consolidate reports suggesting SlpA as the surface receptor for *C. difficile* phage infection, we showed that ΦCD1801 was able to bind its own RT but unable to bind to RT 012 or RT 027 strains. Plasmid-borne expression of the RT 078-SLC (H2/6) in RT 012 and RT 027 strains was able to remedy this phenomenon, thus confirming the notion that SlpA, the major constituent of the S-layer, represents the phage receptor on the surface of *C. difficile*.

## Methods

### Routine growth of *C. difficile* strains

*C. difficile* isolates were routinely grown on brain heart infusion (BHI) medium supplemented with 0.5 % yeast extract, 0.1 % L-cysteine and *C. difficile* selective supplement comprising 250 µg/mL D-cycloserine and 8 µg/ml cefoxitin (Oxoid, USA), referred to as BHIs. Strains were maintained at 37°C under anaerobic conditions in a Don Whitley anaerobic workstation (80% N_2_, 10% CO_2_ and 10% H_2_).

### Isolation of *C. difficile* from stool samples

Novel C. *difficile* isolates used within this study were isolated from patient faecal samples collected at the Queens Medical Centre, Nottingham, UK. Stool samples were homogenised 1:1 with PBS, heat shocked at 80°C for 15 min and centrifuged for 5 min at 1500 *x* g. A 50 μL aliquot of the supernatant was used to inoculate triplicate Cycloserine Cefoxitin Egg Yolk (CCEY) agar plates (LabM, UK) in an anaerobic cabinet and incubated for 48 h. Plates were prepared by autoclaving 48g premixed CCEY in 1L dH2O and adding 40 mL (4%) egg yolk emulsion (Lab M, UK) post-autoclave. Prior to use, the plates were kept under anaerobic conditions for a minimum of 4 h. Putative *C. difficile* isolates were transferred into a 96 microtiter plate containing 200 μL BHIS broth, one per well and up to 20 per patient sample. Microtiter plates were sealed with breathable sterile film and incubated overnight in anaerobic conditions. A separate 96 well microtiter plate contained 180 μL PCR grade H_2_O where a 1:10 dilution was made from the overnight broth cultures. A drop of glycerol was then added to the broth cultures and resealed using fresh breathable sterile film and stored at -80°C. The H_2_O culture mix was covered in breathable sterile film and stored at -20°C for subsequent use as a PCR template for ribotyping. A complete list of strains used in this study is provided in Table S1.

### Ribotyping

Ribotyping of the clinical isolates was performed exactly as described previously [22], following the extraction of DNA from the abovementioned treated stool samples by heating at 95°C for 20 min after initial defrosting. PCR products were visualised using a Qiaexcel using the Qiaexcel DNA High Resolution (Qiagen, Germany) . Band profiles were analysed by eye in the first instance before sending each isolate to the *C. difficile* Reference Network (CDRN) at Leeds Royal Infirmary (Leeds, UK) where for official assignment of strain ribotype.

### Isolation of phages

Sewage samples were obtained from an anaerobic digester at Stoke Bardolph sewage treatment plant in Nottinghamshire, UK. The sewage sample (50 mL) was enriched overnight, anaerobically, with the dry components of BHIs with the addition of 1% taurocholate and MgCl_2_. Subsequently, the enrichment cultures were centrifuged at 10,000 x g for 10 min at 4°C to remove bacteria and debris. The supernatant was filter sterilised (0.22 µM filter, Millipore) and stored at 4°C. Potential RT 078 hosts were selected from isolates obtained at the Queens Medical Centre, Nottingham, UK. Phages were identified through plaque formation and plaques were subsequently purified three times. Lysogens of the isolated phage within the propagating strain CD1801, were isolated using the spot on the plate method as previously described [23]. Lysogens were then confirmed by induction of prophage using Mitomycin C (3 µg/mL) in accordance with established methods [24]. Finally, lysogens were screened for their immunity to further phage infection by means of plaque assay.

### Enumeration of phages

Phages were enumerated using the double agar overlay plaque assay, in accordance with a published protocol [25].

### Transmission electron microscopy

Isolated phage lysates (>10^9^ pfu/mL), were precipitated by 1M ammonium acetate (Sigma Aldrich, USA) as described by Fortier and Moineau (2007), with centrifugation steps at 21,000 x g for 75 min [14]. The precipitated phage particles were stained with 10 µL of 2% uranyl acetate (Sigma Aldrich, USA) for 30s on 200 mesh Formavar carbon coated copper grids, before visualising through transmission electron microscopy (TEM) following an established method [14].

### Extraction of phage genomic DNA

Phage genomic DNA was extracted from crude phage lysate using a modified phenol/chloroform method [8]. A 2 mL volume of crude phage lysate (∼10^9^ pfu/mL) was mixed with 25 µL MgCl_2_ (1M, Sigma), 0.8 µL DNase I (2000 U/ml, Thermo Fisher Scientific) and 20 µl RNAse A (10 mg/ml, Thermo Fisher Scientific) and incubated at room temperature for 30 min. Subsequently, 80 µL EDTA (0.5 M, Thermo Fisher Scientific), 5 µL Proteinase K (20 mg/mL, Qiagen) and 100 µl 10% SDS (Thermo Fisher Scientific) were added to the phage-MgCl_2_ mixture and incubated at 55 °C for 1 h. The resulting liquid was aliquoted into 4 phase lock tubes (Quanta Biosciences) and extracted 3 times with an equal volume of phenol:chloroform:isoamylalcohol (25:24:1, Sigma). A final extraction with an equal volume of chloroform (Sigma) was conducted before the DNA was precipitated using 2 volumes 100% ethanol and 0.1 volumes sodium acetate (Sigma) and incubated on ice for 5 min. All centrifugation steps were performed at 13,000 rpm for 5 min. The precipitated DNA was centrifuged at 13,000 rpm for 10 min and resulting pellet washed with 1 ml 70% ethanol. The centrifugation step was repeated and the pellet air-dried before the DNA was dissolved in 10 mM Tris-HCl (pH 8.5, Qiagen) at 65 °C for 20 min. Eluted DNA was pooled, quantified using NanoDrop Lite spectrophotometer (Thermo Scientific) and stored at 4 °C before sequencing.

### Sequencing and annotation of genomes

Full genome sequencing of *C. difficile* strain CD2315 and purified phages was conducted by DeepSeq (University of Nottingham) using an Illumina MiSeq platform. For CD2315, paired reads were aligned to the archetypal genome sequence of strain M120. For the phage genomes, raw sequencing reads were assembled into a single contig using *de novo* assembly function within CLC Genomics Workbench 9.5.3 (Qiagen). Artemis software [26] was used to identify putative open reading frames (ORFs). Manual genome annotation was completed using NCBI BLASTp, UniProt and pfam databases to assign putative protein functions. ORFs were manually trimmed to the correct start codon based on the presence of ribosome binding sites and promoter sequences.

### Determination of phage host range testing

Standard plaque assay was used to determine the host range of the isolated phage using a ∼10^9^ pfu/mL stock according to established methods [25]. Efficiency of plating was determined for each indicator strain by comparison of the phage titer using the propagating strain against the phage titer using the indicator strain (EOP = phage titer of propagating strain ÷ phage titer of indicator strain). Tested strains are listed in Table S1. PHAge Search Tool Enhanced Release (PHASTER) was used to identify prophage regions within the genome of resistant isolates [18]. Putative repressor proteins were aligned using The European Molecular Biology Open Software Suite (EMBOSS) pairwise alignment tool [16].

### S-layer receptor testing

pRPF185 plasmids expressing the hybrid RT 078 S-layer cassette (H2/6) (pJAK002) and the individual SLCs 2 (pJAK023) and 6 (pJAK018), under the control of an anhydrotetracycline-inducible promoter, were obtained from Robert Fagan (University of Sheffield, UK) [27]. pRPF185 was conjugated into *C. difficile 630* exactly as described previously using *E. coli* CA434 as a conjugal donor strain [28]. A 1% inoculum of an overnight culture of the transformed strain was transferred to 20 mL pre-reduced BHIs broth and incubated for 4 h before being induced with anhydrotetracycline (Sigma Aldrich, USA) to a final concentration of 500 ng/mL in a 20 mL culture for 1h under anaerobic conditions. To detect phage binding in the presence and absence of the RT 078 S-layer cassette, a binding assay was conducted. Therein, 20 mL of induced cultures were harvested by centrifugation and the resulting cell pellet re-suspended in 10 µL of phage (10^4^ pfu/ml). This was incubated for 15 min under anaerobic conditions to allow the phage to bind before being re-suspended in 1 mL BHIs broth. A final centrifugation step was completed to remove the bacterial cells. The number of phage particles in the supernatant that had not bound to cells was enumerated using plaque assay as mentioned above, using CD1801 as an indicator. A substantial reduction in phage titer from the infection phage titer is indicative of phage binding. *C. difficile* 630 and 1801 were used as negative and positive binding controls, respectively.

## Acknowledgements

We thank Rob Fagan (University of Sheffield) for his kind donation of SLC-expressing derivatives of pRPF185 and his advice regarding the S-layer. We would also like to thank Erasmus students Arlen-Celina Lücke (University of Hanover) and Renske van Esveld (Leiden University) for their contribution to laboratory work. This work was supported by a Medical Research Council Industrial CASE (MRC; grant No. MR/K017829/1) with Phico Therapeutics Ltd and by the NIHR Nottingham BRC (Reference no. BRC-1215-20003). The views expressed are those of the authors and not necessarily those of the funders.

N.P.M conceived the study and all experimental work on phage isolation and characterisations was undertaken by M.J.W. M.M.L. isolated and characterised all Nottingham-derived *C. difficile* 078 strains, M.J.W. and T.W.B. undertook all genome analysis and annotation and drafted the manuscript. All authors reviewed, edited, and approved the final version of the manuscript.

We declare no conflicts of interest.

## Supplementary information

**Table S1:**
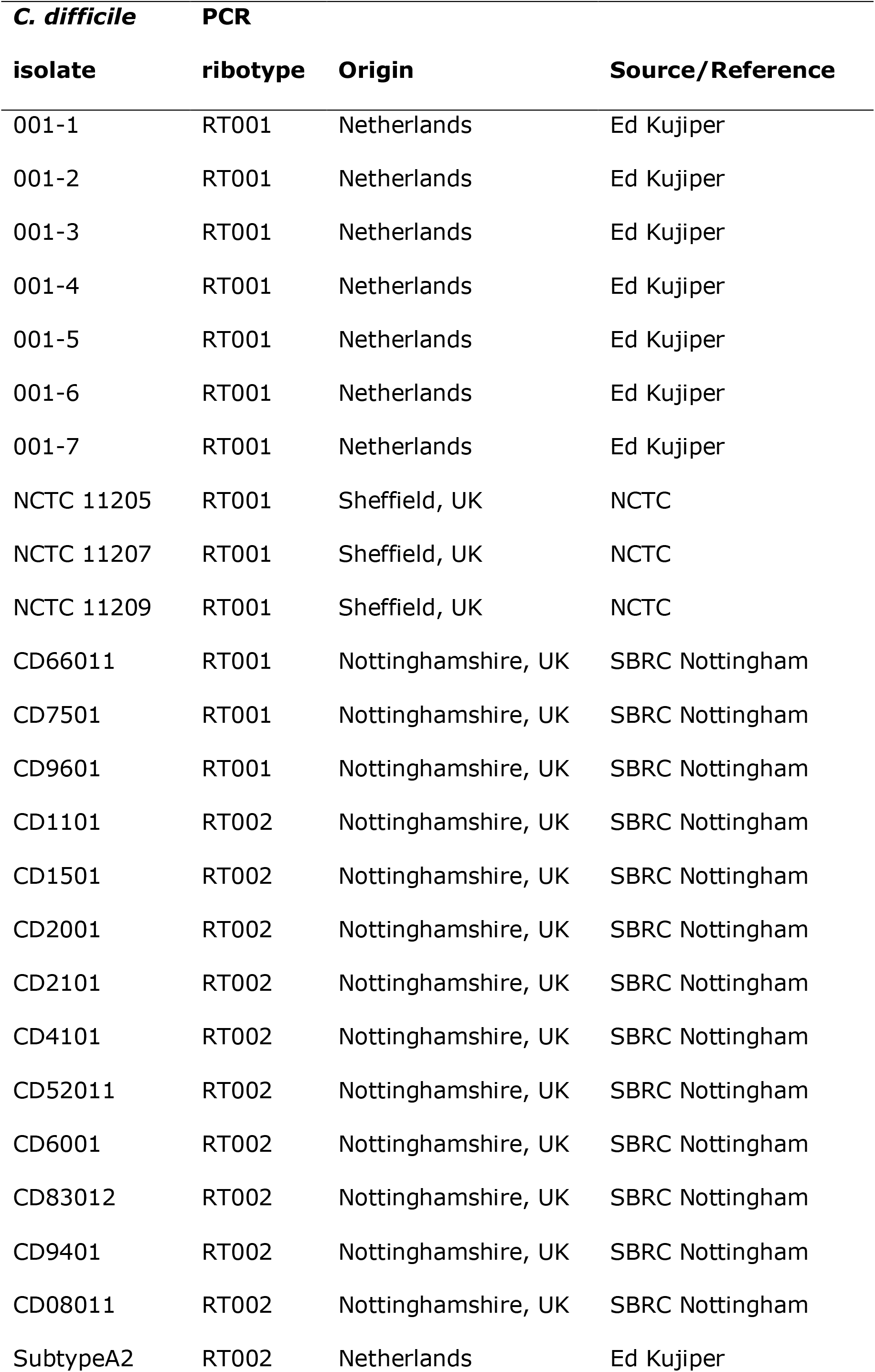
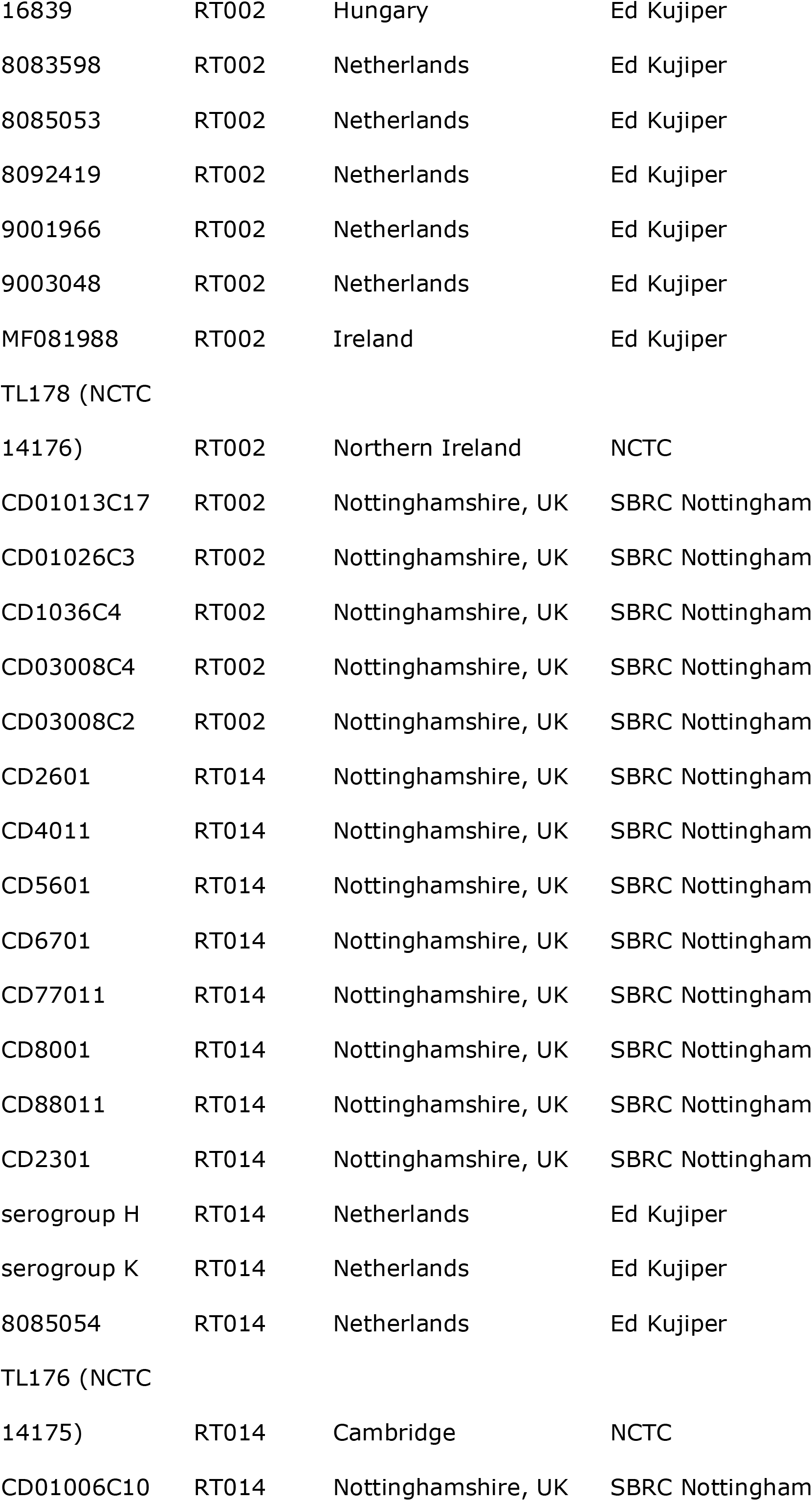
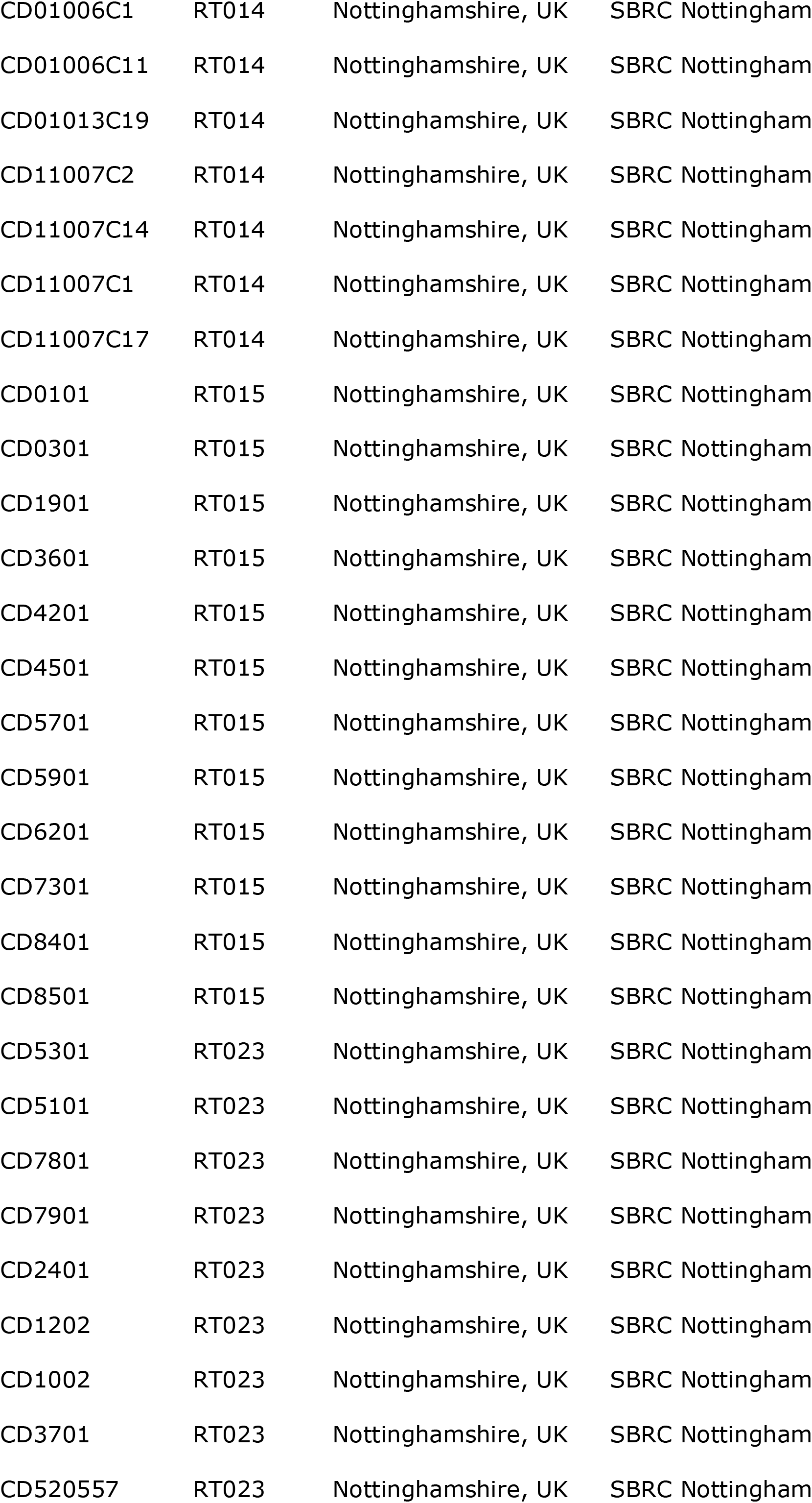
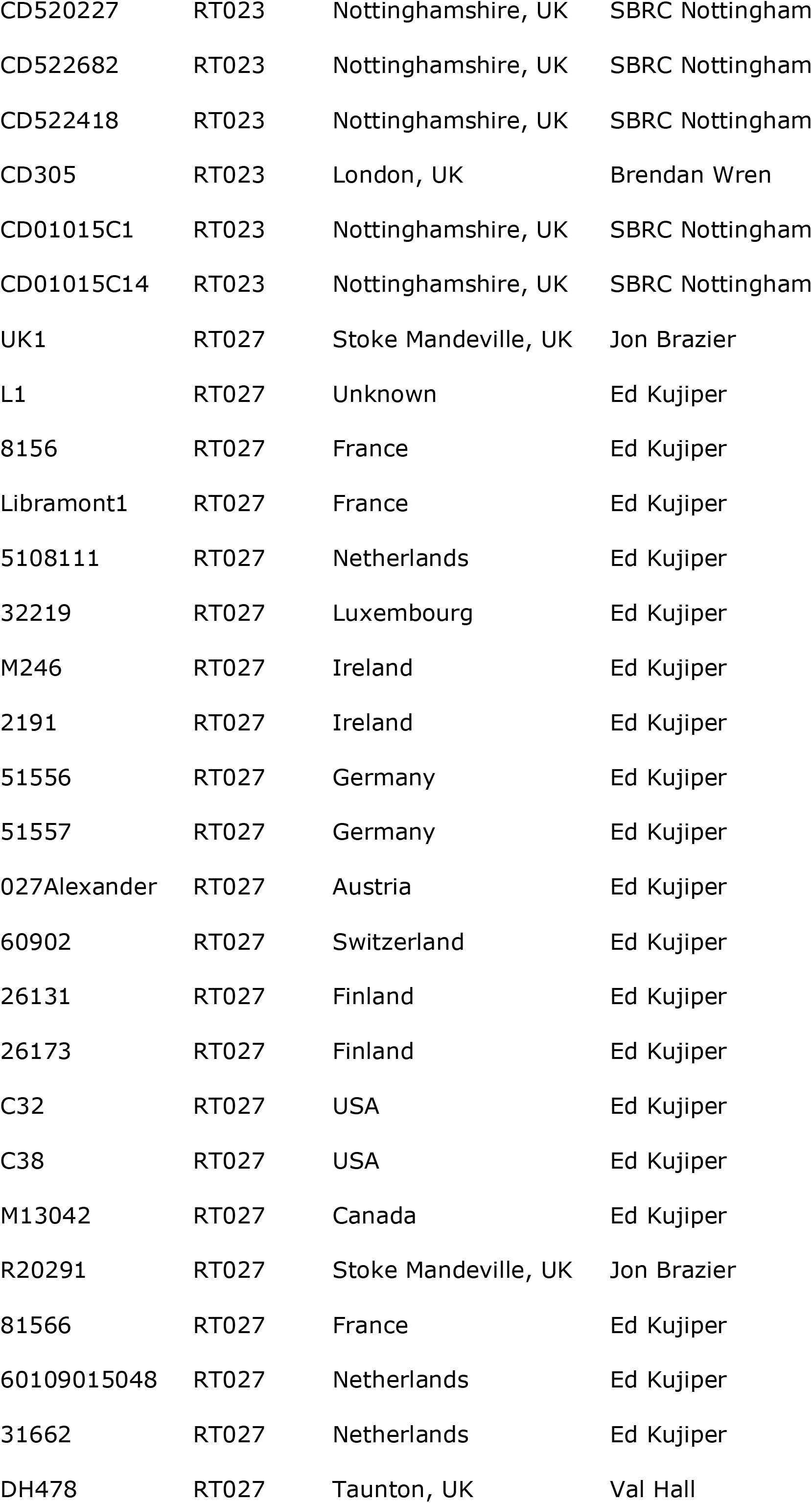
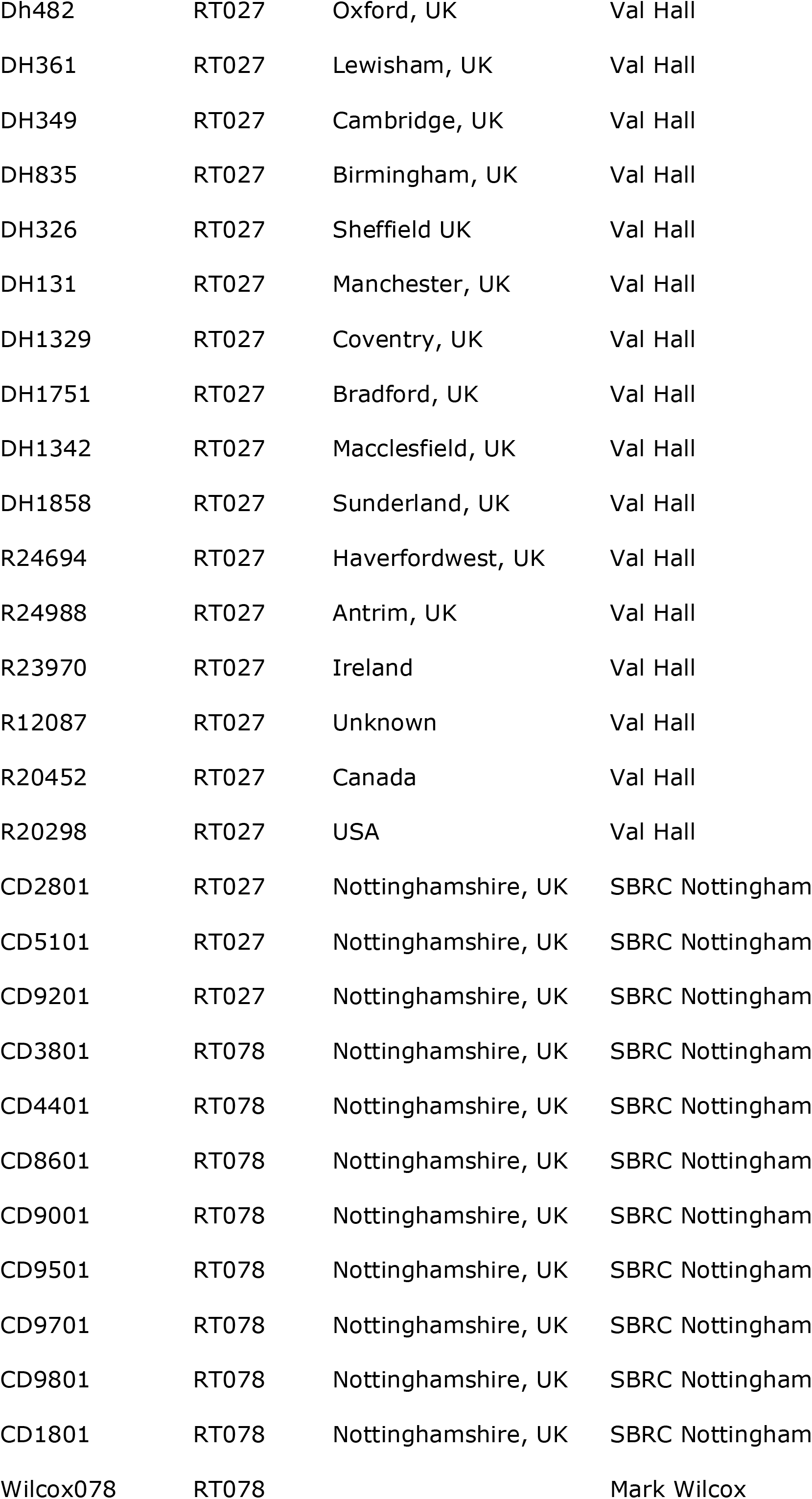
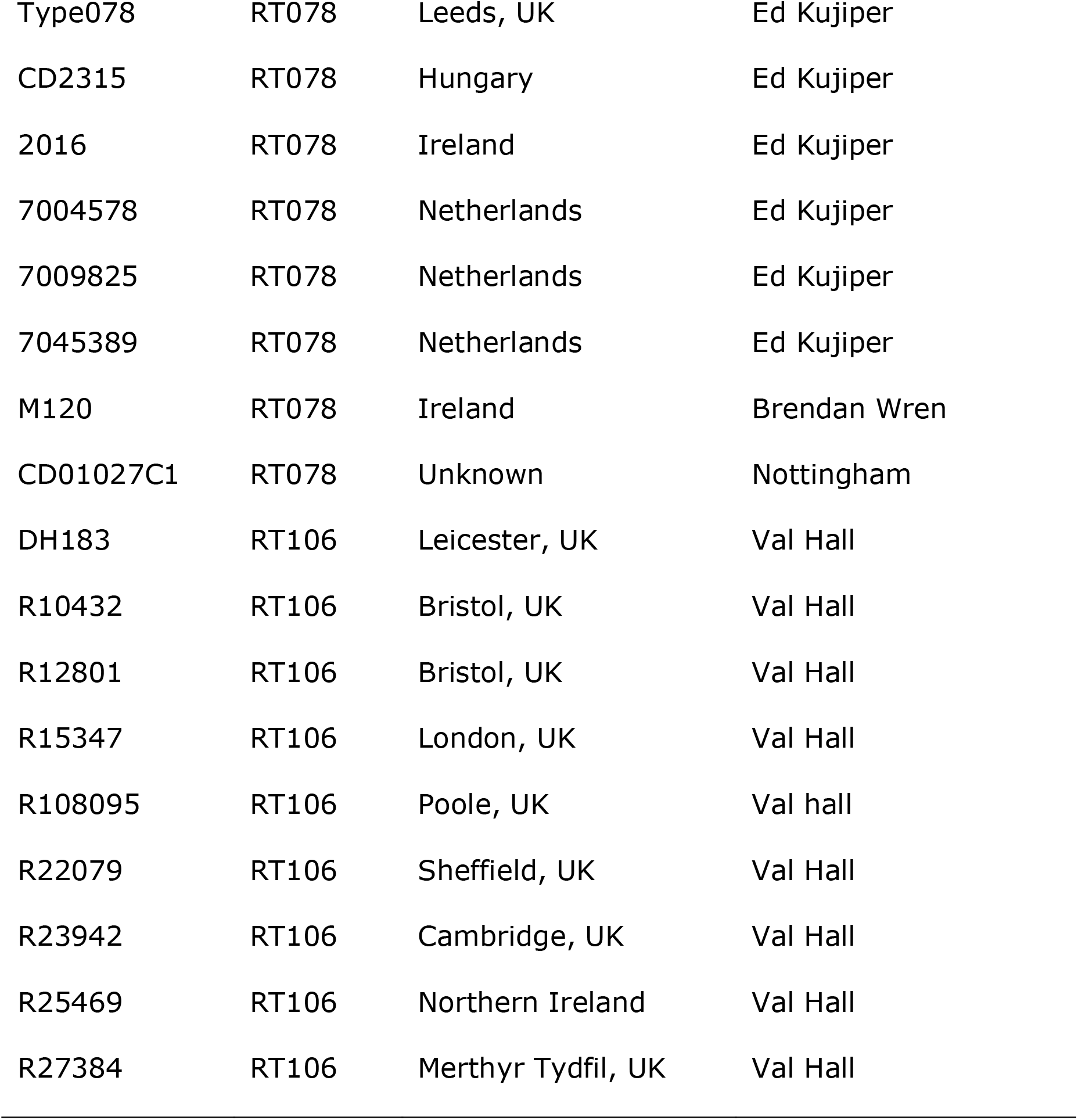
*C. difficile* isolates used in this study.

**Table S2:**
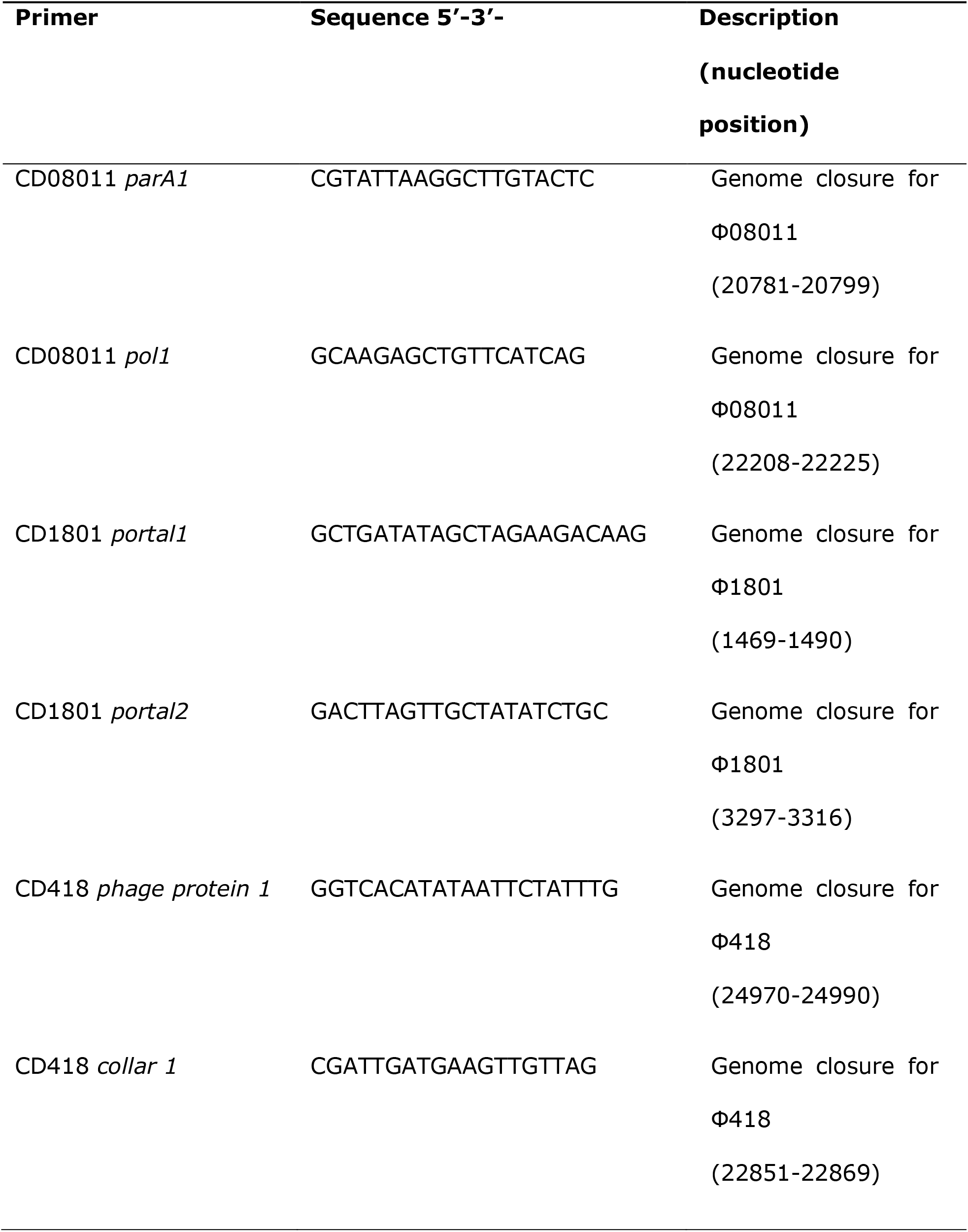
List of primers for the closure of phage genomes.

**Figure S1:**
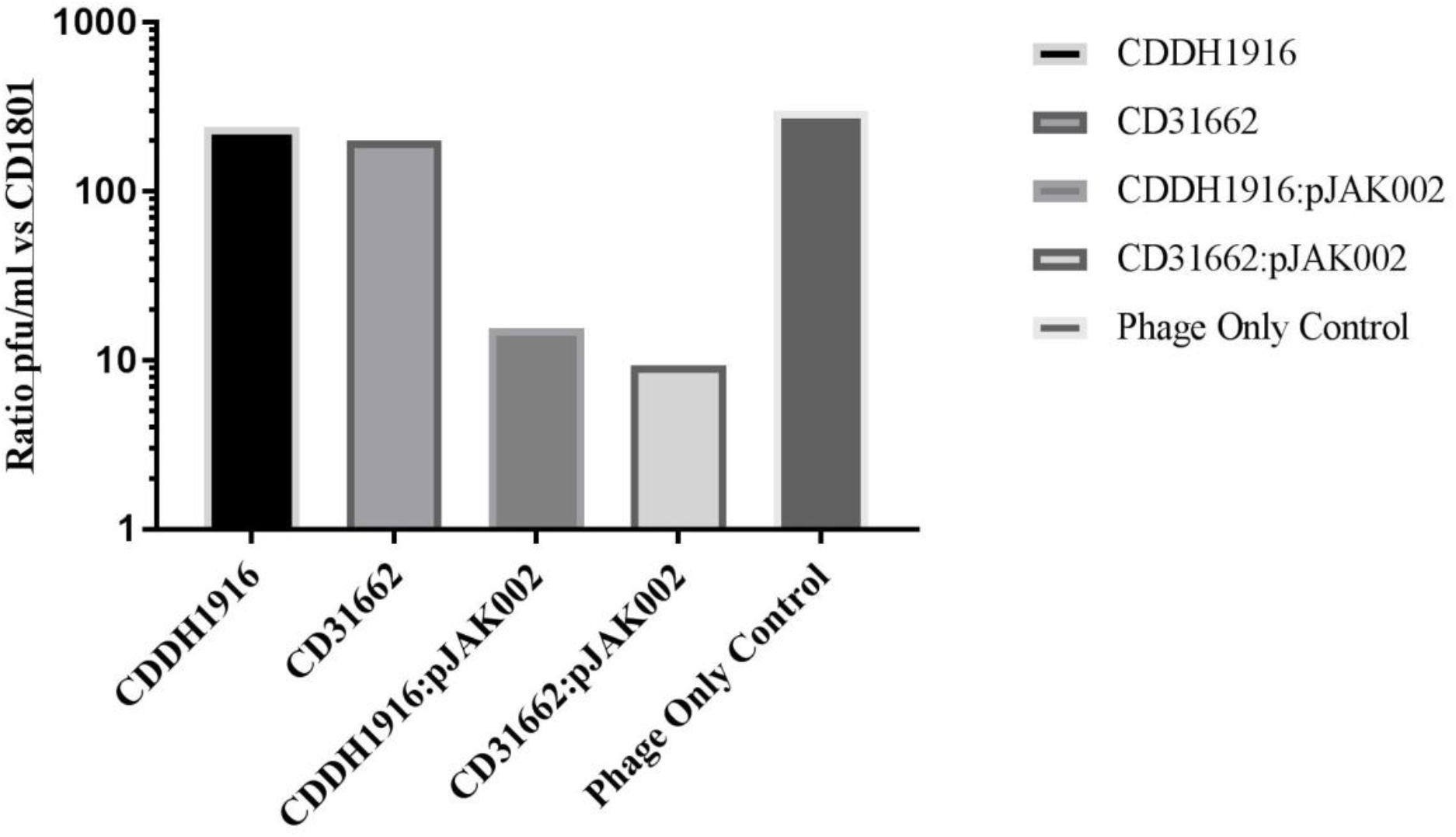
Phage binding is SLCT-dependent in RT 027. The capacity for CDDH1916 and CD31662 to bind to ΦCD1801 was assessed by means of a phage binding assay with wild-type parental strains and those harnessing plasmid-borne SLC H2/6 (pJK002) under the control of a tetracycline inducible promoter. Following co-incubation with wild-type or plasmid-bearing RT 027 strains, the titer of free ΦCD1801 particles was assessed through a plaque assay using the propagating strain CD18101 as an indicator. Data represent the mean of a single biological experiment and as such, statistical confidence cannot be obtained.

## References

1. Lawson PA, Citron DM, Tyrrell KL, Finegold SM. 2016. Reclassification of *Clostridium difficile* as *Clostridioides difficile* (Hall and O’Toole 1935) Prevot 1938. Anaerobe 40:95–9.

2. Lessa FC, Mu Y, Bamberg WM, Beldavs ZG, G.K. Dumyati, J.R. Dunn, M.M. Farley, S.M. Holzbauer, J.I. Meek, and E.C. Phipps. 2015. Burden of *Clostridium difficile* infection in the United States. N Engl J Med 372(24):2369–70.

3. Theriot CM, Young VB. 2014. Microbial and metabolic interactions between the gastrointestinal tract and *Clostridium difficile* infection. Gut Microbes 5(1):86–95.

4. Higa JT, Kelly CP. 2014. New drugs and strategies for management of *Clostridium difficile* colitis. J Intensive Care Med. 29(4):190–9.

5. Chang JY, Antonopoulos DA, Kalra A, Tonelli A, Khalife WT, Schmidt TM, Young VB. 2008. Decreased diversity of the fecal Microbiome in recurrent *Clostridium difficile*-associated diarrhea. J Infect Dis 197(3):435-8.

6. Ross A, Ward S, Hyman P. 2016. More Is Better: Selecting for Broad Host Range Bacteriophages. Front Microbiol 7:1352–1352.

7. Krutova M, Zouharova M, Matejkova J, Tkadlec J, Krejci J, Faldyna M, Nyc O, Bernardy J. 2018. The emergence of *Clostridium difficile* PCR ribotype 078 in piglets in the Czech Republic clusters with *Clostridium difficile* PCR ribotype 078 isolates from Germany, Japan and Taiwan. Int J Med Microbiol 308(7):770–775.

8. Nale JY, Spencer J, Hargreaves KR, Buckley AM, Trzepiński P, Douce GR, Clokie MRJ. 2016. Bacteriophage Combinations Significantly Reduce *Clostridium difficile* Growth *In Vitro* and Proliferation *In Vivo*. Antimicrob Agents Chemother 60(2):968–981.

9. Kirk JA, Gebhart D, Buckley AM, Lok S, Scholl D, Douce GR, Govoni GR Fagan RP+. 2017. New class of precision antimicrobials redefines role of *Clostridium difficile* S-layer in virulence and viability. Sci Trans Med 9(406):eaah6813.

10. Phothichaisri W, Ounjai P, Phetruen T, Janvilisri T, Khunrae P, Singhakaew S, Wangroongsarb P, Chankhamhaengdecha S. 2018. Characterization of Bacteriophages Infecting Clinical Isolates of *Clostridium difficile*. Front Microbiol 9:1701–1701.

11. Wu Y-C, Lee JJ, Tsai BY, Liu YF, Chen CM, Tien N, Tsai PJ, Chen TH. 2016. Potentially hypervirulent *Clostridium difficile* PCR ribotype 078 lineage isolates in pigs and possible implications for humans in Taiwan. Int J Med Microbiol 306(2):115–122.

12. Goorhuis A, Bakker D, Corver J, Debast SB, Harmanus C, Notermans DW, Bergwerff AA, Dekker FW, Kuijper EJ. 2008. Emergence of *Clostridium difficile* infection due to a new hypervirulent strain, polymerase chain reaction ribotype 078. Clin Infect Dis 47(9):1162–70.

13. O’Connor JR, Johnson S, Gerding DN. 2009. *Clostridium difficile* Infection Caused by the Epidemic BI/NAP1/027 Strain. Gastroenterol 136(6):1913–1924.

14. Fortier L-C, Moineau S. 2007. Morphological and Genetic Diversity of Temperate Phages in *Clostridium difficile*. Appl Environ Microbiol 73(22):7358–7366.

15. Rashid SJ, Barylski J, Hargreaves KR, Millard AA, Vinner GK, Clokie MRJ. 2016. Two Novel Myoviruses from the North of Iraq Reveal Insights into *Clostridium difficile* Phage Diversity and Biology. Viruses 8(11):310.

16. Rice P, Longden I, Bleasby A. 2000. EMBOSS: the European Molecular Biology Open Software Suite. Trends Genet 16(6):276–7.

17. Mayer MJ, Narbad A, Gasson MJ. 2008. Molecular characterization of a *Clostridium difficile* bacteriophage and its cloned biologically active endolysin. J Bacteriol 190(20):6734–40.

18. Arndt D, Grant JR, Marcu A, Sajed T, Pon A, Liang Y, Wishart DS. 2016. PHASTER: a better, faster version of the PHAST phage search tool. Nucl Acids Res 44(W1):W16–21.

19. Kirk JA, Banerji O, Fagan RP. 2017. Characteristics of the *Clostridium difficile* cell envelope and its importance in therapeutics. Microb Biotechnol 10(1):76–90.

20. Fagan RP, Albesa-Jove D, Qazi O, Svergun DI, Brown KA, Fairweather NF. 2009. Structural insights into the molecular organization of the S-layer from *Clostridium difficile*. Mol Microbiol 71(5):1308-22.

21. Ge P, Scholl D, Leiman PG, Yu X, Miller JF, Zhou ZH. 2015. Atomic structures of a bactericidal contractile nanotube in its pre-and postcontraction states. Nat Struct Mol Biol 22(5):377–82.

22. O’Neill G, Ogunsola F, Brazier J, Duerden B. 1996. Modification of a PCR Ribotyping Method for Application as a Routine Typing Scheme for *Clostridium difficile*. Anaerobe. 12(4):205–9.

23. Govind R, Vediyappan G, Rolfe RD, Dupuy B, Fralick JA. 2009. Bacteriophage-mediated toxin gene regulation in *Clostridium difficile*. J Virol 83(23):12037–12045.

24. Sell TL, Schaberg DR, Fekety FR. 1983. Bacteriophage and bacteriocin typing scheme for *Clostridium difficile*. J Clin Microbiol 17(6):1148–1152.

25. Clokie MRJ, Kropinski A. (2009). Bacteriophages. Methods and Protocols: Isolation, Characterization, and Interactions, Vol. 1, eds M. R. J. Clokie and A. M. Kropinski (Totowa, NJ: Humana Press).

26. Rutherford K, Parkhill J, Crook J, Horsnell T, Rice P, Rajandream MA, Barrell B. 2000. Artemis: sequence visualization and annotation. Bioinformatics 16(10):944–5.

27. Fagan RP, Fairweather NF. 2011. *Clostridium difficile* has two parallel and essential Sec secretion systems. J Biol Chem 286(31):27483–93.

28. Cartman ST, Minton NP. 2010. A mariner-Based Transposon System for In Vivo Random Mutagenesis of *Clostridium difficile*. Appl Environ Microbiol 76(4):1103–1109.

